# Identification of potential blood-derived extracellular vesicles biomarkers to diagnose and predict distant metastases in ER+ breast cancer patients

**DOI:** 10.1101/202291

**Authors:** Zia Hamidi, Eduardo Tejero, Ronny Schmidt, Rod Tucker, Ana Pedro

## Abstract

The objective of this study was to identify potential extracellular vesicles (EV) which might serve as biomarkers for predictive and diagnostic purposes in metastatic breast cancer. Plasma samples from 7 different metastatic and non-metastatic ER+ (estrogen receptor -positive) breast cancer patients were collected, EV were isolated and their protein content analyzed by mass spectrometry and FunRich analysis. In this study we found several putative plasma EV biomarkers for metastatic ER+ breast cancer prediction and diagnosis, such as serum amyloid A1, known to promote widespread metastasis in a breast cancer animal model.

## Introduction

EVs are membrane surrounded structures released by different cell types that are involved in cellular communication. EVs are involved in several of the steps of metastasis including the detachment of tumor cells through epithelial to mesenchymal transition (EMT) and allowing the physical dissemination to distant organs. Tumor-derived EVs are also critical components for preparing the tumor microenvironment because they enable tumor cells to escape from the immunological surveillance **(1)** and help in the setting of a pre-metastatic niche for the engraftment of detached cancer cells **(2)**. For these reasons, EVs are potential therapeutic and diagnostic targets in cancer and EV-derived biomarkers maybe useful for predicting future metastatic development **(3)**. Adjuvant therapy is used to reduce the risk of breast cancer (BC) progression to metastases. However, most of the patients would survive without this treatment as their cancers do not metastasize **(4)**. Here we describe several possible candidate plasma EV-derived biomarkers that could potentially be used to diagnose metastatic ER+ breast cancer and predict distant metastases in early ER+ breast cancer either by using protein-microarrays or flow cytometry or ELISA. In addition, the presence of such biomarkers could help identify those patients who would benefit from adjuvant therapy or of biphosphonates treatment to prevent bone metastases or to detect earlier brain metastases by detecting the presence of circulating EVs containing ceruloplasmin.

## Material and Methods

### Human samples

Plasma samples from 4 control patients (2 adult women and men) (Lyden lab, Weill Cornell Medicine, New York, USA) and samples CF37, CF5, CF110, CF1, CF25, CF27 and CF (Champalimaud Foundation) 33 (CCC, Breast Unit, Lisbon, Portugal). These samples were collected at Champalimaud Clinical Centre (CCC), Portugal RECI/BIM-ONC/0201/2012 (FCT, Portugal), as part of a study on the role of tumor-derived microvesicles and bone marrow progenitor cells as diagnostic and prognostic biomarkers in advanced (ABC) and inflammatory breast cancers (IBC) patients”. The proteomic profiles of the samples were submitted to Vesiclepedia (http://microvesicles.org/index.html).

### EV purification and analysis

EV purification and analysis were performed at the Lyden lab (WCM) accordingly to Andre et al. 2016 **(5)**. Briefly, cell supernatant or plasma samples were centrifuged for 20 min. at 20,000xg. The supernatant was collected and again centrifuged for 70 min at 100,000xg. The supernant was discarded and the pellets were ressuspend in PBS (phosphate-buffered saline) and centrifuged once more for 70 min at 100,000xg. The supernatant was again discarded and the pellets ressuspend in 100 µl of PBS.

### Proteomics and proteomic analysis

Proteomic analysis was performed at the Rockefeller University, Proteomics Resource Center. Identification of exosomal proteins was performed using reversed phase high pressure liquid chromatography-mass spectrometry (HPLC-MS). Samples were denatured at 90°C, reduced with 10mM DTT at 51°C for 1 h and alkylated with 50mM iodoacetamide at 25°C for 45 min. Proteins were digested with trypsin (Promega, Madison, WI, V5280) overnight at 25°C. Tryptic peptides were concentrated by vacuum centrifugation and desalted using in-house made C18STAGE Tips prior to mass spectrometric analysis.

Samples were loaded by an Eksigent AS2 autosampler onto a 75 µm fused silica capillary column packed with 11cm of C18 reverse phase resin (5 µm particles, 200Å pore size; Magic C18AQ; Michrom BioResources Inc., Auburn, CA, USA). Peptides were resolved on a 180 minute 1-100% buffer B gradient (buffer A = 0.1 mol/l acetic acid, Buffer B = 70% acetonitrile in 0.1 mol/l acetic acid) at a flow rate of 200nl/min. The HPLC (1200 series; Agilent, Santa Rosa, CA, USA) was coupled to a mass spectrometer (LTQ-Orbitrap;ThermoFisher Scientific, Carlsbad, CA, USA) with a resolution of 30,000 for full MS followed by seven datadependent MS/MS analyses. Collision-induced dissociation (CID) was used for peptide fragmentation. Each sample was analyzed at least twice. All MS data were analyzed with Proteome Discoverer software (version 1.2; Thermo Fisher Scientific, San Jose, CA) to search against human and mouse UniProt databases. The peptides were constrained to be tryptic and up to 2 missed cleavages were allowed. Carbamidomethylation of cysteine was specified as a fixed and oxidation of methionine as a variable modification. The precursor ion tolerance was set to 25 ppm, and fragment ion mass tolerance to 0.8 Da. Search results were analyzed individually and data for replicates were combined and evaluated. Proteomic analysis was performed with the help of FunRich Program (http://www.funrich.org/). Only proteins with Mascot scores approximately 90 or bigger than 90 were considered.

## Results and discussion

Details of the EVs isolated from BC patients plasma are shown in Table 1. The extraction process also precipitates contaminants such as lipoproteins and immunocomplexes (IC) **(6)**. However, all of these samples including controls contain transmembrane or lipid-bound extracellular proteins, cytosolic proteins with membrane or receptor binding activity and intracellular proteins associated with other compartments rather than plasma membrane or endosomes as described in Lotvall et al., 2014 **(7)** what confirms they contain EVs. We performed FunRich Venn Diagram analysis with CF37, CF5, CF110, CF1, CF27, CF25 and CF33 EV proteomes to identify specific plasma EV protein biomarkers related to metastatic breast cancer and with progression of early breast cancer to metastases (original mass spectrometry files corresponding to these samples can be found at www.dropbox.com, C@d@vez888, provisonally, and at Vesiclepedia). For protein identification, the protein is typically cleaved by trypsin and the resulting peptides analyzed by MALDI MS/MS or by nano-LC ESI MS/MS. The mass spectrometric data contain a list of the accurate peptide masses and peptide fragment masses. These are searched against sequence databases containing known protein amino acid sequences using the Mascot database search software. The Mascot Score is a statistical score for how well the experimental data match the database sequence. The Mascot score for a protein is the summed score for the individual peptides, e.g. peptide masses and peptide fragment ion masses, for all peptides matching a given protein. For a positive protein identification, the mascot score has to be above the 95% confidence level. For a database search in the nrbd protein database from NCBI containing 18 million known protein sequence, the 95% confidence level is around a Score of 90 **(8)**.

**Table 1:**
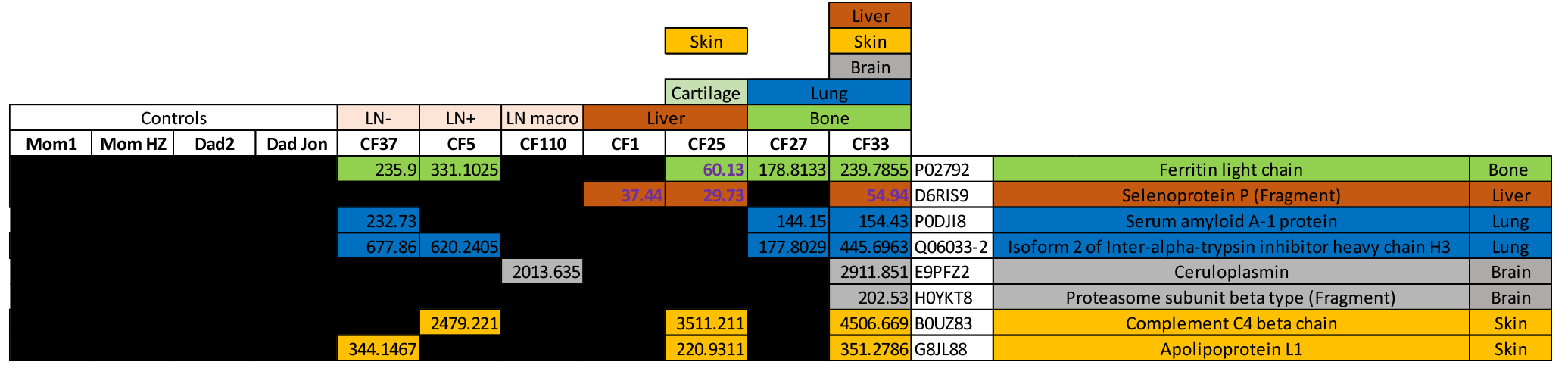
Plasma EV-derived candidate biomarkers for metastatic ER+ breast cancer and metastasis prediction. Mom 1, mom HZ, Dad2 and Dad Jon are plasma control samples derived from 2 women and two men respectively. CF37 is a sample derived from a LN- patient (T2N0), CF5 from a LN+ patient (T2N1), CF110 a sample derived from a patient with macrometastasis in the LNs (T3N1), CF1 a sample derived from a patient with liver metastasis (T3N2M+), CF25 is from a patient with pubic synphesis, liver and skin metastasis (T4N1), CF27 from a patient with lung and bone metastasis (T4N2M1) and CF33 from a patient with liver, skin, bone, lung and brain metastasis (T4). Also, represented the Mascot scores for each protein in each sample. Marked in blue are non-significant Mascot scores.

Only proteins with Mascot scores approximately 90 or greater than 90 were considered.

### Bone marrow alterations

We found ferritin light chain in CF27 and CF33 samples both derived from patients with bone metastases and with Mascot scores of 178.8 and 239.7 respectively (Table 1). Additionally, ferritin light chain was present in samples CF37 (LN-negative) and CF5 (LN-positive) but not in control samples. This suggests that EV-derived ferritin light chain may represent a potential marker for bone marrow alterations in breast cancer patients and eventually bone metastasis.

### Lung metastases

Two putative biomarkers for liver metastatases are serum amyloid A-1 protein and isoform 2 of inter-alpha-trypsin inhibitor heavy chain H3 (table 1). Both proteins were present in the samples with lung metastases (CF27 and CF33) but absent in both controls and samples without lung metastases. In addition, these two proteins are also present in the CF37 sample, which may indicate that this patient could develop lung metastases as serum amyloid A-1 is known to induce metastasis in animal models of breast cancer **(9)**. However, the inter-alpha-trypsin inhibitor heavy chain H3 was shown to reduce metastatic spreading **(10)**. These apparently contradictory findings require further work to more clearly define the role of both proteins in metastatic disease.

### Brain metastases

Once again, two putative candidate biomarkers, ceruloplasmin and proteasome subunit beta type (fragment) both of which were absent in controls were present in the sample from the patient with brain metastases (CF33). Brain metastasis are naturally chemoresistant and indeed ceruloplasmin, a ferroxidase enzyme, is among the six candidate serum proteins that were found to predict responses to neoadjuvant chemotherapy in advanced breast cancers **(11)**. Preclinical studies also demonstrate that proteasome inhibition potentiates the activity of other cancer therapeutics, in part by downregulating chemoresistance pathways **(12)**. Because these are both present it suggests that chemoresistance pathways are set up providing a good soil for the onset of brain metastases.

### Skin metastases

Finally, potential biomarker candidates for skin metastasis are complement C4 beta chain and apolipoprotein L1 which are also absent in control samples

Though these initial observations may represent potential biomarkers for the distal metastatic disease, a recognised limitation of the present study is the small sample size (7 ER+ breast cancer patients with different metastatic and non-metastatic profiles) and only 4 EV control samples. Nevertheless, a strength of our study is that samples were drawn from those with confirmed metastatic disease at the different sites and so are likely to be representative patients. Further work in a larger cohort of patients is necessary to confirm these initial findings.

## Conclusion

These preliminary finding potentially suggest a set of EV-derived biomarkers such as serum amyloid A1 that could be used to identify both early breast cancer and whether the disease has metastasized to other sites. If confirmed in a large patient cohort, the biomarkers could be isolated and incorporated into a non-invasive test that could be used for the detection of patients with metastatic ER+ breast cancer and identify the sites to which the cancer has metastasized. This would help clinicians decide which patients will benefit of adjuvant therapy.

## Ethics approval and consent to participate

This study was approved by an Ethics Review Board at Champalimaud Foundation (CF), Portugal. All patients that participated in this study provided their informed consent.

## Availability of data and material

The mass spectrometry analysis results from all the patient samples used in this study are stored provissonaly in the in following dropbox: www.dropbox.com, C@d@vez888. The data will be available at Vesiclepedia.

## Acknowledgements

We are very thankful to Leonid Schneider, independent science journalist at https://forbetterscience.com/ for his help concerning research integrity and publishing matters.

